# VBCG: 20 validated bacterial core genes for phylogenomic analysis with high fidelity and resolution

**DOI:** 10.1101/2023.06.13.544823

**Authors:** Renmao Tian, Behzad Imanian

## Abstract

**Background:** Phylogenomic analysis has become an inseparable part of studies of bacterial diversity and evolution, and many different bacterial core genes have been collated and used for phylogenomic tree reconstruction. However, these genes have been selected based on their presence and single-copy ratio in all bacterial genomes, leaving out the gene’s ‘phylogenetic fidelity’ unexamined.

**Results:** From 30,522 complete genomes covering 11,262 species, we examined 148 bacterial core genes that have been previously used for phylogenomic analysis. In addition to the gene presence and single-copy rations, we evaluated the gene’s phylogenetic fidelity by comparing each gene’s phylogeny with its corresponding *16S rRNA* gene tree. Out of the 148 bacterial genes, 20 validated bacterial core genes (VBCG) were selected as the core gene set with the highest bacterial phylogenetic fidelity. Moreover, the smaller gene set resulted in a higher species presence in the tree than larger gene sets. Using *Escherichia* and *Salmonella* as examples of prominent bacterial foodborne pathogens, we demonstrated that the 20 VBCG produced phylogenies with higher fidelity and resolution at species and strain levels while *16S rRNA* gene tree alone could not.

**Conclusion:** The 20 validated core gene set improves the fidelity and speed of phylogenomic analysis. Among other uses, this tool improves our ability to explore the evolution, typing and tracking of bacterial strains, such as human pathogens. We have developed a Python pipeline and a desktop graphic app (available on GitHub) for users to perform phylogenomic analysis with high fidelity and resolution.

## Background

Bacterial pangenomes consist of the core genome, a set of genes present in all genomes within a clade, and the accessory genome, a set of genes found in one or more but not all members of the clade [1]. While the core genome of a clade can be, and often is, different from that of another clade, it invariably contains highly conserved genes essential for the vital cellular functions of the members of that clade.

The core genome reflects the vertical accumulation of mutations on the conserved set of genes, and thus it can provide phylogenetic signals for assigning bacterial strains to their corresponding populations [2]. Over the past decade, there has been a significant increase in research on bacterial core genes and their applications in phylogenomic analysis [3–6]. Bacterial core genes are widely used for phylogenomic analysis due to a) their high level of conservation; and b) their widespread distribution among bacterial genomes. With the increased availability of massive amounts of genomic data and improved phylogenetic methods in recent years, phylogenomic analysis has become a popular approach to study bacterial evolution, diversity and ecology. Phylogenomic analysis using core genes [7] has also been shown to be more accurate and robust than the traditional phylogenetic methods that use a single gene (e.g. *16S rRNA*) or multiple genes (e.g. MLST).

With different number of marker genes, often within a core gene set, different teams of researchers have developed tools and pipelines to simplify and streamline core gene phylogenomic analysis. For example, Wu et al. used 31 predefined marker genes to build a phylogenomic pipeline named AMPHORA [8,9]. These 31 marker genes included those involved in glycolysis, DNA replication, translation etc. Also, Markus et al. developed a bioinformatic pipeline called bcgTree that used 107 essential single-copy core genes, found in a large number of bacteria, to reconstruct their phylogenetic history [6]. This pipeline automatically extracted these genes using hidden Markov models and performs a partitioned maximum-likelihood analysis. Another group presented an up-to-date bacterial core gene set, named UBCG, which consisted of single-copy, homologous genes present in most known bacterial species [4]. The 92 UBCG gene set was selected from 1,429 species covering 28 phyla with the criteria of 90% presence ratio and 90% single-copy ratio. The method was successfully used to infer phylogenomic relationships in *Escherichia* and related taxa, and it can be used at any taxonomic level for Bacteria. Three years later, the same group updated their bacterial core genes to UBCG2 [5], which included 81 genes from 3,508 species spanning 43 phyla with the updated criteria of 95% presence ratio and 95% single-copy ratio.

All the previous studies have utilized the gene presence and single-copy ratios to screen for the conserved, single-copy genes to build their phylogenies. However, this has led to an obvious shortcoming: the selected genes were not examined for their fidelity in reconstructing the phylogenetic tree, and thus they may carry incongruent evolutionary signals. A core genome phylogeny is reconstructed, like other multiple-gene trees, based on concatenated sequences of a set of selected genes. The inclusion of a gene, whose tree shows discordance with the phylogenies of the others genes in the gene set lowers the evolutionary signal and thus the accuracy of topology and resolution of the resulting phylogeny. We, thus, propose that in order to select the genes for a core genome to reconstruct an accurate phylogeny, in addition to high presence and single-copy ratios, the genes should be examined for phylogenetic fidelity, a measure of congruity of the phylogenies of the genes within a gene set, as well. Here, we used the 16s rRNA gene trees to evaluate and compare the phylogenetic fidelity of the candidate core genes and identified 20 high fidelity genes for bacterial phylogenomic analysis. We have developed a pipeline, VBCG, which uses the core gene set to build phylogenomic trees automatically with input of genomic sequences.

## Implementation

### Test genomes and candidate core genes

Genome information of prokaryotes was accessed Through the National Center for Biotechnology Information (NCBI) (https://www.ncbi.nlm.nih.gov/genome/browse#!/prokaryotes/) on Oct 11, 2022. In total, 30,552 finished bacterial genomes covering 11,262 species were selected and the protein sequences and rRNA genes were downloaded using a custom Python script with multiple processing. The rRNA files were then filtered to remove those with multiple *16S rRNA* genes that were less than 99% identical. For those with multiple copies that were > 99% identical, the first copy was used as a representative. The *16S rRNA* genes were then clustered using CD-HIT (version 4.8.1) [10] with a similarity threshold of 0.99 and an alignment coverage of 0.9 (-c 0.99 -G 0 -aL 0.9). The representative sequences (5506 sequences) were then randomly divided into 100 groups for further analysis.

We used a total of 148 genes collected from UBCG [4], UBCG2 [5] bac120 [11] and bcgTree [6] as candidate genes, which have been shown to have high presence and single-copy ratios in bacterial genomes. To acquire the HMM model files, PGAP HMM files and a PFAM file were downloaded from an NCBI website (https://ftp.ncbi.nlm.nih.gov/hmm/current/) and the European Molecular Biology Laboratory’s European Bioinformatics Institute (EMBL-EBI) website (https://ftp.ebi.ac.uk/pub/databases/Pfam/current_release/), respectively, and the corresponding HMM models were retrieved using a custom Python script.

To annotate the candidate core genes, hmmscan of the package HMMER [12,13] was used to search all the protein sequences of the 5506 representative genomes against the acquired HMM models (with the trusted score cutoffs, --cut_tc), in parallel using Python package multiprocessing. The output file was parsed to assign the core genes to the proteins.

### Ubiquity and uniqueness of the core genes

The ubiquity and uniqueness of the core genes were evaluated by calculating the presence ratio and single-copy ratio of the core genes in the 5506 representative genomes. The presence ratio was defined by the number of genomes with each core gene divided by the total number of genomes. The single-copy ratio was defined by the number of genomes with only one copy of each core gene divided by total number of genomes. Only the core genes with both presence ratio and single-copy ratio > 95%, respectively, were used for further analysis.

### Fidelity test for the candidate core genes

For each candidate core gene, all the proteins were divided into 50 groups corresponding to the grouping of *16S rRNA* genes. For each group, a phylogenetic tree was reconstructed for the *16S rRNA* gene and the core gene, respectively. The sequences were first aligned using MUSCLE (version 3.8.1551) [14]. The multiple sequence alignment was then trimmed to remove terminal gaps. The trimmed alignment was then filtered to select conserved blocks using Gblocks (version 0.91b) [15] with the minimum length of a block as three (-b4=3) and the maximum number of allowed gap positions as half (-b5=h). The resulted alignment was then converted into format Phylip and was then fed to FastTree (version 2.1.10) [16,17] for phylogenetic tree reconstruction. The models gtr + gamma and lg + gamma were used for the *16S rRNA* gene and core gene protein sequences, respectively, with a bootstrap number of 100. The output trees in Newick format of *16S rRNA* gene and core genes of each group were compared using Dendropy (version 4.5.2) [18] calculating the Robinson Foulds (RF) distance [19]. The 20 core genes with the highest fidelity, also ensuring that > 80% of the genomes include the whole gene set, were chosen as the validated bacterial core gene (VBCG) set.

### Comparison of multiple gene sets in fidelity

The 50 groups of genomes were used to compare the fidelity of the gene set VBCG with other gene sets. For each group, the core genes were aligned using MUSCLE and the terminal gaps were removed from the multiple alignments. Gblocks was then used to select conserved blocks. The processed alignments of the core genes were then concatenated, removing any taxa with > 1 genes missing. The concatenated alignments were converted into Phylip files and fed to FastTree for phylogenetic tree reconstruction. The trees were then compared to the corresponding *16S rRNA* gene trees by calculating the RF distance.

## Results

### Ribosomal protein-encoding genes have high ubiquity and uniqueness in the Kingdom Bacteria

In total 30,522 rRNA gene and their corresponding protein sequences were downloaded from RefSeq database. After filtering those with multiple divergent copies, 29,781 sequences of *16S rRNA* genes were remained. After clustering the sequences at 99%, 5,506 representative genes were observed, which were then randomly divided into 50 groups (**Figure 1**).

**Figure 1.**
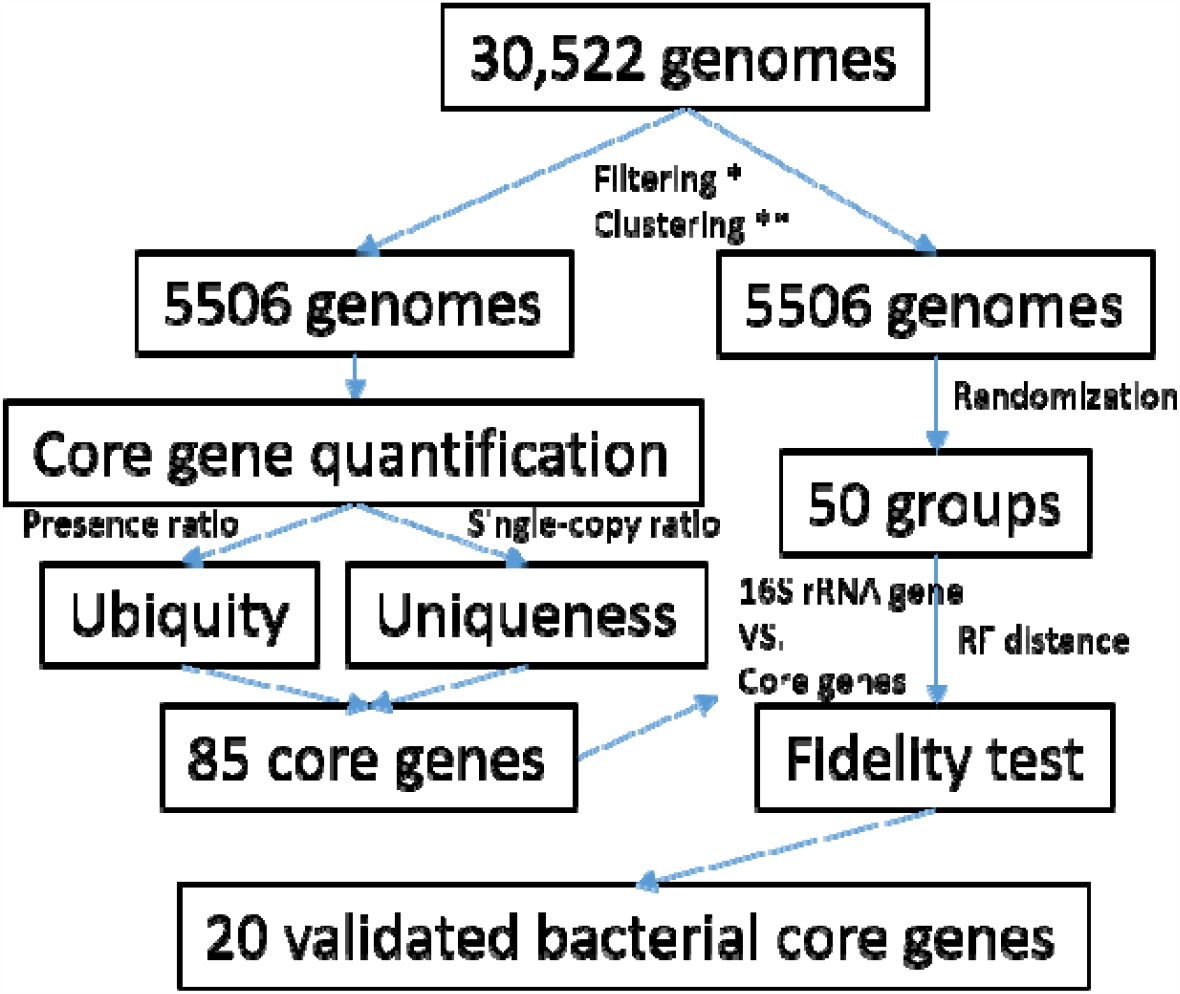
The workflow of the data analysis in this study. The filtering step indicated by an asterisk removes genomes with > 2 copies of *16S rRNA* genes that are < 99% identical. The clustering step indicated by two asterisks clusters *16S rRNA* genes using a cutoff of 99%

The presence and single-copy ratios of the 148 core genes in the 5506 representative genomes were calculated (**Figure 2A**). Interestingly, the 47 ribosomal proteins had much higher ubiquity (median presence ratios: 98.9%) and uniqueness (median single-copy ratios: 98.7%) than those of the other core genes (95.7% and 94.2%). Among the 50 proteins with top presence ratios, 39 were ribosomal proteins. We then selected the core genes (**Figure 2B, Table S1**) with both presence and single-copy ratios > 95% for the fidelity and resolution tests (85 genes).

**Figure 2.**
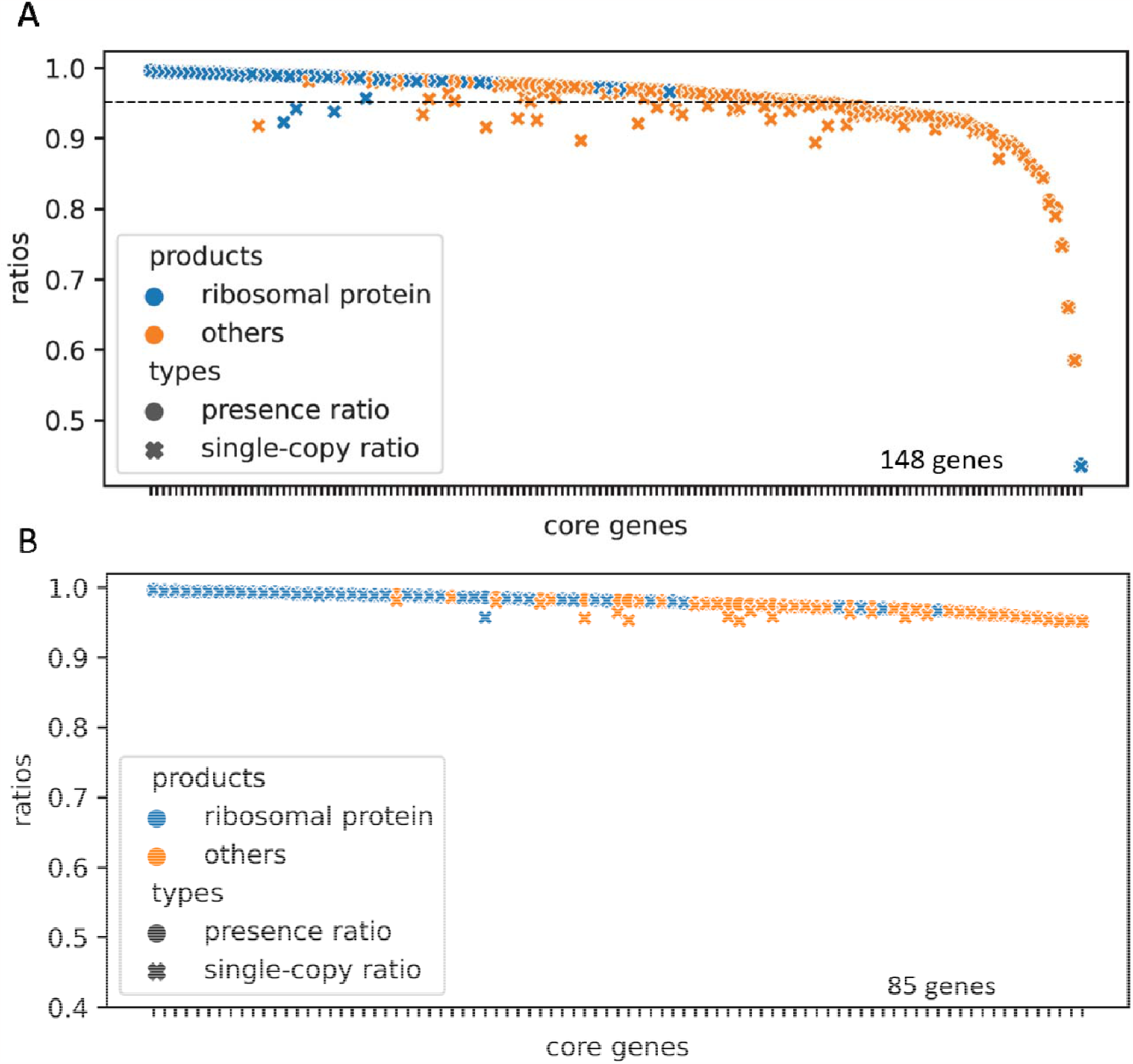
The presence and single-copy ratios of the candidate core genes in the Kingdom Bacteria. In the total core genes (**A**), the 47 ribosomal proteins had much higher ubiquity (median presence ratios: 98.9%) and uniqueness (median single-copy ratios: 98.7%) than those of the other core genes (95.7% and 94.2%). The dashed line demarcates the 95% ratio. In total 85 core genes (**B**) with both presence and single-copy ratios > 95% were retained for the fidelity and resolution tests.

### The validated core gene tree showed the highest concordance with the 16S rRNA gene phylogeny

The 5,506 representative genes were randomly divided into 50 groups (∼110 taxa in each group, **Figure 1**). The fidelity test compared the trees of the 50 groups for each of the 85 core genes with the corresponding *16S rRNA* gene trees based on Robinson-Foulds (RF) distance. The tests ranked the fidelity for each of the core genes (**Figure 3A**). The core gene phylogenies with the closest and farthest distances to their corresponding *16S rRNA* gene trees ranged from 115.9 ± 14.8 RF and 173.8 ± 9.1 RF, respectively, with the maximum being 50% higher the minimum. We then chose the top 20 genes with the highest fidelity (with lowest RF distances) as our candidate validated bacterial core gene (VBCG) set. A close examination of these 20 genes revealed that they were mostly involved in transcription (e.g. RNA polymerase) and translation (e.g. ribosomal proteins, translation initiation, elongation and termination).

**Figure 3.**
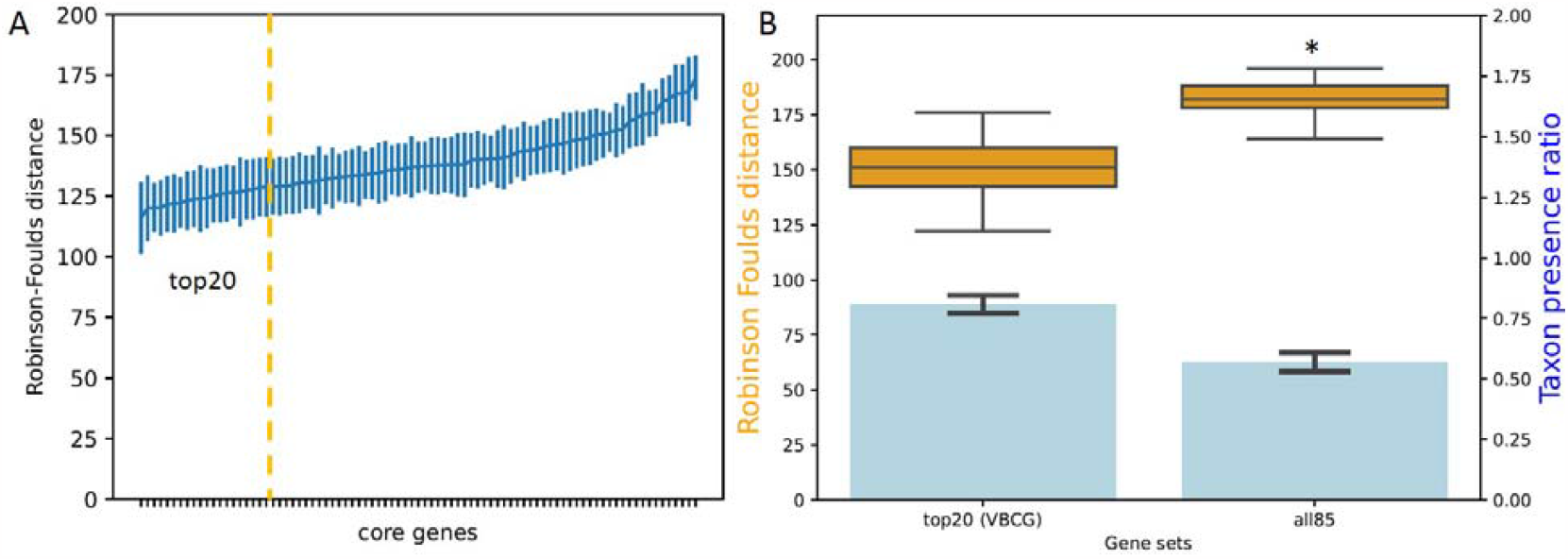
The fidelity test of the validated core genes. The fidelity tests (A): The RF distances of the core genes trees from the corresponding *16S rRNA* gene trees. Each data point indicates the means and standard deviations of 50 trees (110 taxa each). The yellow dashed line indicates the top 20 genes with the highest fidelities in the left. The fidelity test and comparisons for phylogenies of multiple gene sets (B): The RF distances (brown boxes) of the trees based on the concatenated genes (top 20 gene set and all 85 gene set) from the corresponding *16S rRNA* gene trees of the 50 groups of genomes. The asterisk indicate significant difference from the top 20 gene set. The blue bars and right side y axis show the taxon presence ratio of each gene set, which indicates the percentage of taxa with all members of the gene sets.

We then compared the phylogenetic fidelity of the top 20 gene (VBCG) set and that of all the 85 genes, by concatenating the genes and calculating the RF distances of the resulting trees from the corresponding *16S rRNA* gene trees of the 50 groups of genomes. As a result, the VBCG gene set had significantly lower RF distance from the corresponding *16S rRNA* gene trees (Paired Samples T test, *p* < 1e-5, **Figure 3B**) than those of the 85 gene set, which was 20% higher.

### The VBCG gene set resulted in less species loss in the tree compared to that of all genes

The rationale behind selecting the top 20 genes is that these genes yield an acceptable species loss rate in the phylogenetic tree. Because all the core genes had to have a presence ratio between 95% and 100%, only certain taxa will have all the genes of a given gene set to be concatenated for phylogenetic tree reconstruction. One advantage of the VBCG set over the all 85 gene set is the higher percentage of the taxa that can be included in the tree reconstruction. Of all the 50 groups, averagely 80.7% taxa in each group (**Figure 3B**) had all the VBCG genes present for phylogenetic reconstruction, but this number fell to 57.0% for the all 85 gene set. Our results clearly demonstrate that the inclusion of more genes would result in the loss of taxa as shown in the tree containing all the genes.

### The VBCG gene set phylogeny had better resolution than that of the 16S rRNA gene

We compared the discriminative capability of VBCG set and the *16S rRNA* gene in tree reconstruction on the Genus *Escherichia* with an outgroup species from the genus *Salmonella*, both having member species that are commonly implicated in foodborne disease outbreaks. Our results showed that the *16S rRNA* gene tree had polytomies and could hardly discriminate between the species of *Eschericha* mainly because the *16s rRNA* genes in the main group had a very high sequence similarity (> 99.0% pairwise identities) (**Figure 4A**). The *16s rRNA* tree also suffered from misclassification of the *E. coli* Str. IMT2125, which was mistakenly positioned outside the *Eschericha* genus. Various strains of the species *E. coli*, and *E. fergusonii* grouped together and were not well separated by species. Some of them are at the same branch, implying that they had not diverged. However, the VBCG tree showed a well-defined grouping by species, with different species nicely separated in different clades. The *E. coli* Str. IMT2125 was correctly positioned inside the genus. Even within each species clade, the strains were well separated with different branch lengths. In the VBCG tree, *E. coli* Str. 2020CK-00188 had branched within the *E. fergusonii* clade. While we cannot completely dismiss that this might be a mistake within the VBCG tree that begs further investigations, the likely explanation is that *E. coli* Str. 2020CK-00188 has been misclassified and it is not an *E. coli* species at all. Overall, our results indicated that a phylogeny based on the 20 VBCG genes resulted in higher resolutions (even at the strain level) compared with the corresponding tree based on *16s rRNA* genes.

**Figure 4.**
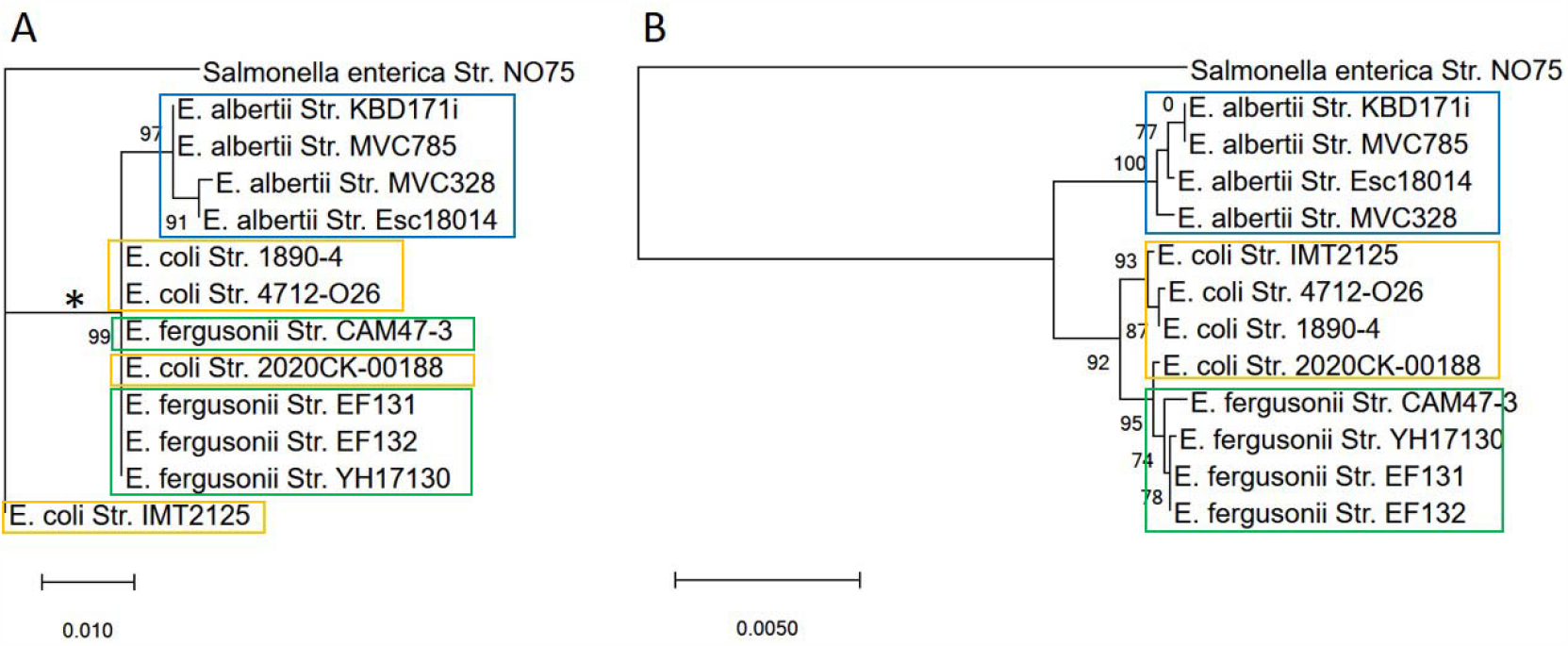
Comparison between *16S rRNA* gene tree (A) and the VBCG tree (B) of concatenated core genes. The tree of the VBCG genes were reconstructed using the concatenated 20 genes of each genome. The asterisk in the *16S rRNA* gene tree indicates the main group of *Escherichia* species with pairwise identities > 99.0%.

### A pipeline and a desktop app for reconstructing phylogenomic trees automatically

We have built a pipeline to reconstruct phylogenomic trees with the input of assembled genomic sequences (**Figure S1**). The pipeline first predicts the gene and protein sequences of the input genomes using Prodigal. The protein sequences are then used to identify the 20 VBCG genes with HMMER. Subsequently, the resulting VBCG genes of all the genomes are retrieved and each gene from all taxa is placed in its own single file. The genes are aligned using Muscle and the terminal gaps are removed from the multiple alignments. Gblocks is then used to select conserved blocks. The processed alignments of the core genes are then concatenated, removing any taxa with > 1 missing gene. The concatenated alignments are converted into Phylip files and fed into FastTree or RAxML for phylogenetic tree reconstruction.

The pipeline is written in Python 3.9, with the following dependencies: Bio >= 1.5.3, Muscle == 3.8, Pandas, Prodigal and HMMER. The command line-based pipeline can be run in Linux operating systems. We have also built a desktop graphic user interface (GUI) app (**Figure S2**) for the users who have no access to a Linux system and a high-performance computer (HPC). The desktop GUI app can be run in Windows and macOS systems.

## Discussion

With the increasing availability of genomic sequences in recent years, phylogenomic analysis has become a powerful critical tool for studying bacterial diversity and evolution. In the past, many marker genes and various core gene sets have been used for phylogenetic tree reconstruction [4,5,8,9,20,21]. However, these genes have been selected based only on their presence and single-copy ratios in all bacterial genomes. The evolution of the genes within a genome is not uniform, and different genes evolve at different rates. Thus, single gene phylogenies, even from the same genome, could be quite different. The inclusion of the genes that are evolving at heterogeneous rates can weaken the phylogenetic signal, introduce noise, produce incorrect topologies, and eventually lower the resolution of the tree. In the previous methods and tools, the gene phylogenetic fidelity has not been considered, compared or tested in order to screen and select the highly concordant marker genes to include in a gene set (e.g. core genome set or its subsets) for phylogenomic analysis. Here, we have identified 20 validated bacterial core genes (VBCGs) that exhibit high fidelity and concordance in phylogeny with their corresponding *16S rRNA* gene trees. Moreover, we demonstrated that the VBCGs could produce superior resolution in discriminating species and strains compared to the widely used *16S rRNA* genes or other core gene sets. We have developed a Python pipeline and a desktop graphic app that enable users to perform phylogenomic analysis using the VBCGs with high fidelity and resolution.

In addition to a better resolution, the 20 VBCG also retains higher number of species and faster run-time in the phylogenomic analysis. Our 85 candidate bacterial core genes based upon the presence and single-copy ratios filtration are consistent with the UBCG2 [5], which contains 81 core genes using the same ratios for filtration. However, our results have shown that the 85 core gene set without fidelity screening resulted in higher RF distances from the corresponding *16s rRNA* trees than the 20 VBCG gene set. In terms of taxon coverage, the 85 core genes resulted in the inclusion of only 57% of all the species in the concatenated sequences, whereas the VBCG gene set included 80% of the species with all the 20 genes. Thus, VBCG can effectively reduce the species loss due to absence of one or more core genes in the phylogenomic tree reconstruction. In addition, the fewer number of core genes in the VBCG tool lowers the running time significantly (up to 4 ×) compared to those with the larger gene sets.

Phylogenomic analysis has emerged as a powerful tool, for example, in tracking the origins of foodborne disease outbreaks. This technique involves the sequencing of whole genomes of bacterial isolates recovered from infected individuals, foods and related environments. These sequences are then used in computational analyses to reconstruct a phylogenomic tree that reveals the evolutionary relationships between these isolates. One of the main benefits of these analyses is their ability to accurately trace the spread of a particular bacterial strain across different geographical locations and over time. Phylogenomic analyses has become an indispensable tool in the hands of epidemiologists as well. A phylogenomic tree of different clinical isolates, for example, can help researchers identify the common ancestors of the pathogens and determine the direction and timeline of the transmission of the disease and the pathogen between individuals within a population. In this study we examined the existing phylogenomic analysis tools and their related core genome gene sets, and we added a phylogenomic fidelity criterion in selecting the core genes. We have developed a new pipeline and tool that relies on a new gene set, the VBCG. The phylogenomic analysis based on the VBGC gene set retains higher number of species, achieves higher speed, and higher resolution than those reliant on other core gene sets or the *16S rRNA* genes as demonstrated in the phylogenomic trees reconstructed for the two common foodborne pathogens, *Escherichia* and *Salmonella*.

## Conclusion

In conclusion, our research used a refined selection process that takes into account gene presence, single-copy ratios, and phylogenetic fidelity, and revealed that the 20 validated bacterial core genes (VBCG) provide high phylogenetic fidelity and resolution for phylogenomic analysis, enhancing our understanding of bacterial diversity and evolution. We have developed a streamlined tool capable of producing more accurate and reliable phylogenies, even at the strain level. Our findings highlight the importance of VBCG in the efficient typing and tracking of bacterial pathogens, a valuable tool in pathogenic studies. We have made our Python pipeline and graphic desktop app available on GitHub to ensure accessibility for other researchers in the field.

## Availability and requirements

Project name: Validated Bacterial Core Genes (VBCG)

Project home page: https://github.com/tianrenmaogithub/vbcg

Operating system(s): Linux, Windows

Programming language: Python

License: GNU GPL v2.0

### List of abbreviations

VBCG: Validated Bacterial Core Genes
NCBI: National Center for Biotechnology Information
EMBL-EBI: European Molecular Biology Laboratory’s European Bioinformatics Institute
RF: Robinson Foulds
GUI: Graphic User Interface
HPC: high-performance computer

## Declarations

### Availability of data and materials

The source code the pipeline VBCG can be accessed on GitHub. In addition, there is a graphic user interface (GUI) version of the pipeline for Windows users (https://github.com/tianrenmaogithub/vbcg).

## Author contributions

B.I. and R.T. conceptualized and designed the project; R.T. conducted the data analysis, code writing and pipeline development. R.T. and B.I. wrote the manuscript. All authors have read and agreed to the published version of the manuscript.

## Acknowledgments

The authors acknowledge the Food and Drug Administration (FDA) of the U.S. Department of Health and Human Services (HHS) for the support.

---

**Table S1.** The 85 core genes with both presence and single-copy ratios > 95%. The presence and single-copy ratios were calculated based on the 5506 representative genomes.

| HMM | Presence ratio | Single-copy ratio | Product |
| --- | --- | --- | --- |
| TIGR01050.1 | 99.6% | 99.6% | 30S ribosomal protein S19 |
| PF00380.22 | 99.5% | 99.4% | Ribosomal protein S9/S16 |
| TIGR01067.1 | 99.5% | 99.5% | 50S ribosomal protein L14 |
| TIGR00981.1 | 99.4% | 99.4% | 30S ribosomal protein S12 |
| PF00281.22 | 99.4% | 99.4% | Ribosomal protein L5 |
| PF00410.22 | 99.4% | 99.4% | Ribosomal protein S8 |
| TIGR01049.1 | 99.4% | 99.3% | 30S ribosomal protein S10 |
| TIGR01164.1 | 99.4% | 99.4% | 50S ribosomal protein L16 |
| PF00276.23 | 99.3% | 99.3% | Ribosomal protein L23 |
| TIGR01029.1 | 99.3% | 99.2% | 30S ribosomal protein S7 |
| TIGR00952.1 | 99.3% | 99.2% | 30S ribosomal protein S15 |
| TIGR00029.2 | 99.3% | 99.2% | 30S ribosomal protein S20 |
| TIGR03631.1 | 99.2% | 99.2% | 30S ribosomal protein S13 |
| TIGR00060.1 | 99.1% | 99.0% | 50S ribosomal protein L18 |
| TIGR01066.1 | 99.1% | 99.0% | 50S ribosomal protein L13 |
| TIGR00855.1 | 99.1% | 98.8% | 50S ribosomal protein L7/L12 |
| TIGR03654.1 | 99.1% | 99.1% | 50S ribosomal protein L6 |
| PF00466.23 | 99.0% | 98.9% | Ribosomal protein L10 |
| TIGR01171.1 | 99.0% | 99.0% | 50S ribosomal protein L2 |
| TIGR01021.1 | 98.9% | 98.9% | 30S ribosomal protein S5 |
| TIGR01032.1 | 98.9% | 98.8% | 50S ribosomal protein L20 |
| TIGR00001.1 | 98.9% | 98.9% | 50S ribosomal protein L35 |
| TIGR02027.1 | 98.9% | 98.1% | DNA-directed RNA polymerase subunit alpha |
| TIGR01071.1 | 98.9% | 98.9% | 50S ribosomal protein L15 |
| TIGR03953.1 | 98.8% | 98.7% | 50S ribosomal protein L4 |
| TIGR03632.1 | 98.8% | 98.7% | 30S ribosomal protein S11 |
| TIGR00062.1 | 98.7% | 98.7% | 50S ribosomal protein L27 |
| TIGR00116.1 | 98.7% | 98.5% | translation elongation factor Ts |
| TIGR00166.1 | 98.6% | 98.5% | 30S ribosomal protein S6 |
| TIGR01169.1 | 98.6% | 98.6% | 50S ribosomal protein L1 |
| TIGR01017.1 | 98.6% | 95.7% | 30S ribosomal protein S4 |
| TIGR00086.1 | 98.5% | 97.9% | SsrA-binding protein |
| TIGR00158.1 | 98.5% | 98.4% | 50S ribosomal protein L9 |
| TIGR03625.1 | 98.4% | 98.4% | 50S ribosomal protein L3 |
| TIGR01079.1 | 98.3% | 98.3% | 50S ribosomal protein L24 |
| TIGR00967.1 | 98.3% | 97.7% | preprotein translocase subunit SecY |
| TIGR00043.2 | 98.3% | 98.2% | rRNA maturation RNase YbeY |
| TIGR00061.1 | 98.3% | 98.2% | 50S ribosomal protein L21 |
| TIGR01632.1 | 98.3% | 98.1% | 50S ribosomal protein L11 |
| TIGR00414.1 | 98.2% | 95.6% | serine--tRNA ligase |
| TIGR01024.1 | 98.2% | 98.2% | 50S ribosomal protein L19 |
| TIGR01030.1 | 98.1% | 98.1% | 50S ribosomal protein L34 |
| TIGR00006.1 | 98.1% | 96.4% | 16S rRNA (cytosine(1402)-N(4))-methyltransferase RsmH |
| TIGR00168.2 | 98.1% | 95.3% | translation initiation factor IF-3 |
| TIGR00755.1 | 98.0% | 98.0% | 16S rRNA (adenine(1518)-N(6)/adenine(1519)-N(6))-dimethyltransferase RsmA |
| TIGR03635.1 | 98.0% | 98.0% | 30S ribosomal protein S17 |
| TIGR00468.1 | 98.0% | 97.9% | phenylalanine--tRNA ligase subunit alpha |
| TIGR01011.1 | 98.0% | 97.9% | 30S ribosomal protein S2 |
| TIGR01044.1 | 97.8% | 97.8% | 50S ribosomal protein L22 |
| TIGR00396.1 | 97.8% | 97.4% | leucine--tRNA ligase |
| TIGR03594.1 | 97.7% | 97.6% | ribosome biogenesis GTPase Der |
| TIGR00496.1 | 97.7% | 97.6% | ribosome recycling factor |
| TIGR00392.1 | 97.6% | 95.8% | isoleucine--tRNA ligase |
| TIGR00442.1 | 97.6% | 95.2% | histidine--tRNA ligase |
| TIGR03723.1 | 97.5% | 96.6% | tRNA (adenosine(37)-N6)-threonylcarbamoyltransferase complex transferase subunit TsaD |
| TIGR00964.1 | 97.5% | 97.4% | preprotein translocase subunit SecE |
| PF00162.22 | 97.5% | 95.8% | Phosphoglycerate kinase |
| TIGR02013.1 | 97.4% | 97.1% | DNA-directed RNA polymerase subunit beta |
| TIGR02729.1 | 97.3% | 97.3% | Obg family GTPase CgtA |
| TIGR00019.1 | 97.3% | 97.3% | peptide chain release factor 1 |
| TIGR00810.1 | 97.2% | 97.0% | preprotein translocase subunit SecG |
| TIGR00092.1 | 97.2% | 97.1% | redox-regulated ATPase YchF |
| TIGR00012.1 | 97.2% | 97.2% | 50S ribosomal protein L29 |
| TIGR01393.1 | 97.2% | 96.3% | translation elongation factor 4 |
| TIGR01009.1 | 97.1% | 97.1% | 30S ribosomal protein S3 |
| TIGR00344.1 | 97.1% | 96.4% | alanine--tRNA ligase |
| TIGR00059.1 | 97.0% | 96.9% | 50S ribosomal protein L17 |
| TIGR02432.1 | 97.0% | 96.9% | tRNA lysidine(34) synthetase TilS |
| TIGR01391.1 | 96.9% | 95.7% | DNA primase |
| TIGR01953.1 | 96.8% | 96.8% | transcription termination factor NusA |
| TIGR02075.1 | 96.7% | 96.1% | UMP kinase |
| TIGR00002.1 | 96.7% | 96.6% | 30S ribosomal protein S16 |
| TIGR00088.1 | 96.6% | 96.6% | tRNA (guanosine(37)-N1)-methyltransferase TrmD |
| TIGR00487.1 | 96.5% | 96.4% | translation initiation factor IF-2 |
| TIGR00959.1 | 96.4% | 96.4% | signal recognition particle protein |
| TIGR00431.1 | 96.2% | 96.1% | tRNA pseudouridine(55) synthase TruB |
| TIGR03263.1 | 96.1% | 96.0% | guanylate kinase |
| TIGR00922.1 | 96.1% | 96.1% | transcription termination/antitermination protein NusG |
| TIGR00020.2 | 95.9% | 95.8% | peptide chain release factor 2 |
| TIGR00064.1 | 95.7% | 95.7% | signal recognition particle-docking protein FtsY |
| TIGR00472.1 | 95.7% | 95.6% | phenylalanine--tRNA ligase subunit beta |
| TIGR00460.1 | 95.5% | 95.3% | methionyl-tRNA formyltransferase |
| TIGR00152.1 | 95.4% | 95.1% | dephospho-CoA kinase |
| TIGR00250.1 | 95.3% | 95.2% | Holliday junction resolvase RuvX |
| TIGR00635.1 | 95.2% | 95.1% | Holliday junction branch migration DNA helicase RuvB |

**Table S2.** The 20 validated bacterial core genes upon the screens based on presence ratio, single copy ratio, and phylogenetic fidelity. The HMM ID of the genes, Robinson-Foulds (RF) distances from the corresponding *16S rRNA* gene trees and the products of the genes were listed.

| Rank | Core genes | RF Mean | RF STD | Products |
| --- | --- | --- | --- | --- |
| 1 | TIGR02013.1 | 115.9 | 14.8 | DNA-directed RNA polymerase subunit beta |
| 2 | TIGR03594.1 | 120.1 | 13.4 | ribosome biogenesis GTPase Der |
| 3 | TIGR01171.1 | 120.3 | 9.9 | 50S ribosomal protein L2 |
| 4 | PF00281.22 | 120.3 | 11.2 | Ribosomal protein L5 |
| 5 | TIGR02027.1 | 121.8 | 11.4 | DNA-directed RNA polymerase subunit alpha |
| 6 | TIGR01011.1 | 122.0 | 12.0 | 30S ribosomal protein S2 |
| 7 | TIGR01009.1 | 122.3 | 10.3 | 30S ribosomal protein S3 |
| 8 | TIGR00959.1 | 123.2 | 11.9 | signal recognition particle protein |
| 9 | TIGR00967.1 | 123.9 | 11.6 | preprotein translocase subunit SecY |
| 10 | TIGR01393.1 | 124.1 | 13.8 | translation elongation factor 4 |
| 11 | TIGR00487.1 | 124.1 | 12.3 | translation initiation factor IF-2 |
| 12 | TIGR01029.1 | 125.3 | 10.9 | 30S ribosomal protein S7 |
| 13 | TIGR01021.1 | 125.8 | 11.4 | 30S ribosomal protein S5 |
| 14 | TIGR00468.1 | 126.5 | 11.7 | phenylalanine--tRNA ligase subunit alpha |
| 15 | TIGR00019.1 | 126.6 | 10.8 | peptide chain release factor 1 |
| 16 | TIGR01169.1 | 126.7 | 14.0 | 50S ribosomal protein L1 |
| 17 | TIGR01953.1 | 127.6 | 12.3 | transcription termination factor NusA |
| 18 | TIGR03625.1 | 127.8 | 12.4 | 50S ribosomal protein L3 |
| 19 | TIGR02729.1 | 128.7 | 11.7 | Obg family GTPase CgtA |
| 20 | TIGR00092.1 | 128.8 | 12.1 | redox-regulated ATPase YchF |

**Figure S1.**
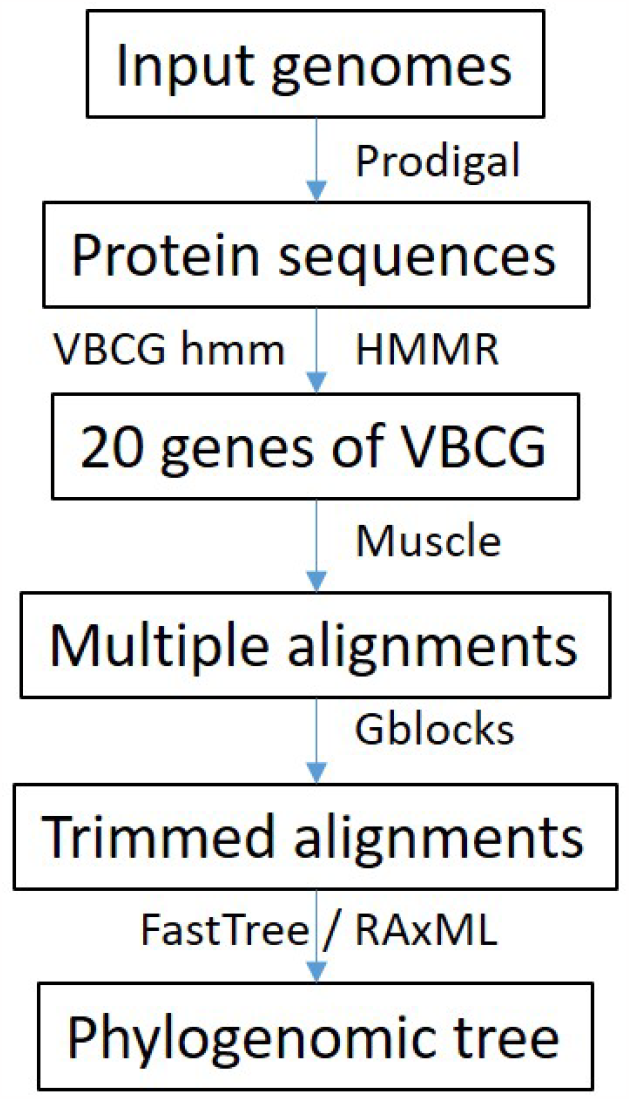
The workflow of the pipeline VBCG. The pipeline begins by predicting gene and protein sequences of the input genomes using Prodigal. Next, the protein sequences are used to identify the 20 VBCG genes with HMMER. The resulting VBCG genes of all the genomes are retrieved and put into separate files for each gene. The genes are aligned using Muscle, and any terminal gaps are removed from the multiple alignments. To select conserved blocks, Gblocks is applied. The processed alignments of the core genes are then concatenated, removing any taxa with more than one gene missing. Finally, the concatenated alignments are converted into Phylip files and fed to FastTree or RAxML for phylogenetic tree reconstruction.

**Figure S2.**
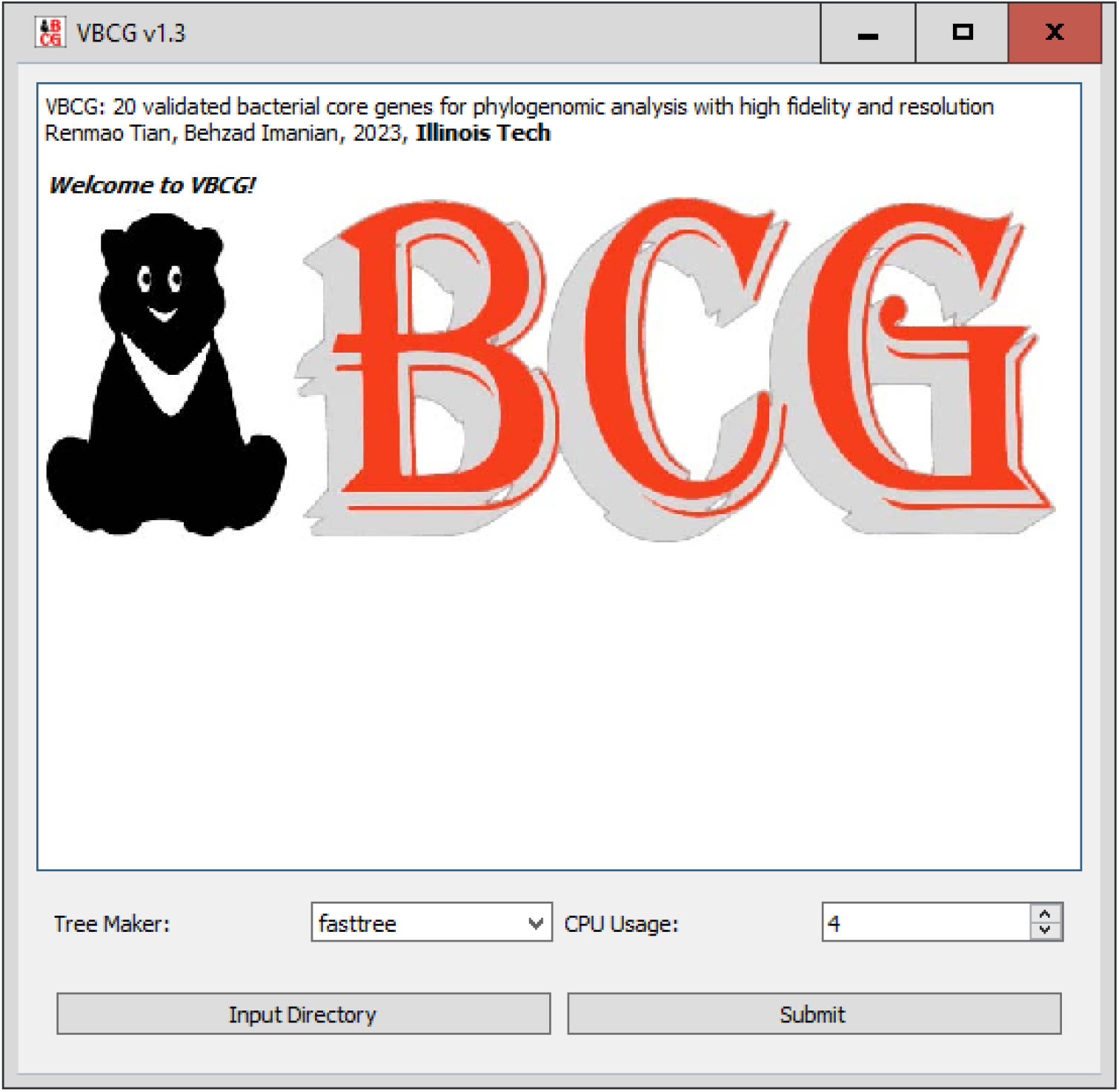
The interface of the VBCG desktop app (Windows version). Users only need to input genomic sequences and set the parameters, and it will identify the 20 VBCG genes with HMMER, and use the concatenated sequence for phylogenomic tree construction.

